# Rectifying AI-generated protein structure ensembles for equilibrium using physics-based computations

**DOI:** 10.64898/2026.03.24.714034

**Authors:** Lisa Otten, Jeremy M. Leung, Lillian T. Chong, Daniel M. Zuckerman

## Abstract

Recently, a number of tools have been released that generate ensembles of protein structures based on artificial intelligence (AI) approaches. Although ensembles generated by the tools differ significantly, we demonstrate a computational path to harmonizing the various outputs under a stationary condition using two complementary physics-based approaches. In the first stage, the AI ensemble is used to seed a weighted ensemble (WE) simulation, promoting relaxation toward the steady state. In the second stage, trajectory segments generated by WE are reweighted to steady state using the recently developed RiteWeight (RW) algorithm. We applied this approach to generate an atomically-detailed equilibrium ensemble of unliganded adenylate kinase conformations, starting from ensembles produced by three AI tools: AFSample2, ESMFlow-PDB (trained from PDB structures), and ESMFlow-MD (trained from molecular dynamics simulation data). Dramatic differences in the AI-generated ensembles are largely erased during the WE-RW process, yielding a consistent description of the equilibrium ensemble for a given force field.

## Introduction

Protein conformational ensembles are now recognized as a central component of modern structural biochemistry, and have arguably eclipsed the single-structure mindset of 20+ years ago. Ensembles provide mechanistic insight into the functional processes that underlie biology, including protein folding,^1,2^ motor protein locomotion,^3,4^ alternating-access active transport,^5,6^ and catalysis coupled to conformational change.^7-9^ Beyond fundamental biology, conformational ensembles are essential to key applications in nanobiology^10^ such as structure-based drug design,^11,12^ the analysis and control of allostery,^13-15^ and protein engineering.^16,17^ In drug design, for example, exploiting “cryptic” allosteric binding sites can lead to more specific or selective drugs than targeting the typically more conserved orthosteric sites.^18,19^ Conformational distributions have been proposed as predictors of cell phenotypes.^20^ Comparisons among ensembles further enable actionable insights into the effects of mutations.^21-23^

Given the biophysical importance of structural ensembles, it is not surprising that the success of “artificial intelligence” (AI) methods, including machine learning, for predicting single protein structures^24-26^ has served to motivate the development of tools for ensemble generation. Our study focuses on three AI-based models: AFSample2 which manipulates the multiple-sequence alignment step in AlphaFold2 to reduce co-evolution information and thus generate a diversity of structure predictions^27^ and two models based on ESMFlow which employs large-language model transformer methodology.^28^ There are two separate ESMFlow models, one based on experimental Protein Data Bank (PDB) structures and the other based on standard molecular dynamics (MD) trajectories in explicit solvent (CHARMM36/TIP3P); although MD trajectories are typically initiated from PDB structures, they should include greater structural diversity based on thermally driven fluctuations.

Our work builds on prior efforts to leverage AI tools for equilibrium sampling.^10,29-37^ Most directly, we build on efforts to pipeline AI-generated ensembles as “seed” structures for subsequent MD simulation and equilibrium analysis by Markov state models.^29,30^ Although very similar in spirit, our approach is methodologically distinct.

Here, we refine AI-generated ensembles using a combination of two existing methods (**Fig. 1**). First, structures from the AI ensemble selected to span the conformational variation are used to initialize weighted ensemble (WE) simulation using the WESTPA software, which is highly optimized for running parallel simulations for systems with complex energy landscapes.^38,39^ WE is unbiased in recapitulating the time-evolution dynamics of a system regardless of binning or other hyper-parameters.^40^ Thus, we expect that after WE simulation the ensemble should be closer to equilibrium than when it started, although computational costs and timescales make it unlikely the system will have reached equilibrium.^41,42^ In the second stage, trajectory segments from WE are used to directly estimate the equilibrium distribution using the RiteWeight algorithm.^43^ RiteWeight uses a self-consistency condition to estimate trajectory segment weights independent of clustering parameters and can be applied to estimate equilibrium or nonequilibrium steady states, a key strength of our approach. We emphasize that RiteWeight is not a standard reweighting protocol based on the importance sampling concept – i.e., ratios of probability distributions and the associated numerical problems – but instead relies on local dynamics, akin to a nearly continuum version of a Markov state model (MSM) ^43^ but without the well-documented initial state bias of MSMs.^44,45^

**Fig. 1.**
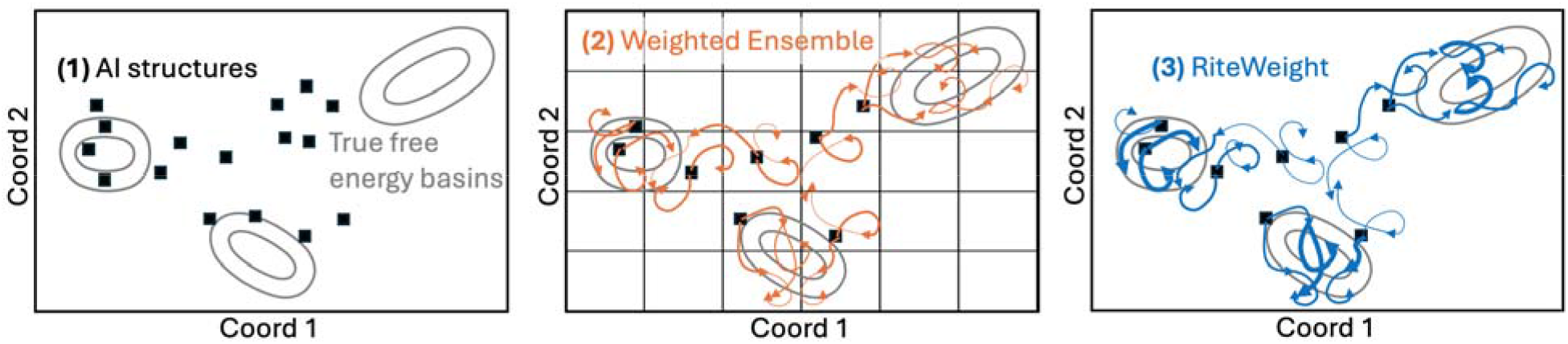
Protein conformational ensembles in three steps. (1) An initial set of protein structures is generated using an AI tool, (2) WE simulation is run from a subset of the AI structures, and (3) RiteWeight reweights the WE trajectories toward equilibrium.

Focusing on the goal of generating an equilibrium conformational ensemble for a protein, our data show that the three chosen AI tools do not provide similar ensembles, implying that further refinement is necessary. Among the tools, it is possible that one of them provides the true equilibrium ensemble while the others are inaccurate, or all of them might be inaccurate. Our data suggest the latter is the case. We emphasize that accuracy, for our purposes, is defined as the Boltzmann-weighted ensemble for a particular force field. In this study, we employ the Amber ff14SB-onlysc protein force field with a generalized Born implicit solvent model (GB-Neck2). No ground-truth ensemble is available, and existing experimental structures come with caveats. X-ray crystal structures will depend on crystallization conditions and crystal packing;^46,47^; NMR structures will depend on solution conditions and the computational refinement algorithm used to determine structures from NMR measurements.^48,49^

Lacking ground truth, we assess equilibrium behavior based on physical principles and self-consistency. Specifically, given that all the AI tools yield different “initial” ensembles, if each can be separately refined into the same final ensemble, that is a strong plausibility argument for success.

## Methods

Our basic pipeline consists of three steps for a given sequence, with full details given below: (1) Generate ensembles of protein structures using the AI tools without modification and compute a joint space of principal components (PCs) using the aggregated ensembles. (2) Subsample each AI ensemble individually using a grid in PC space, and run WE simulation using the same grid separately starting from each AI ensemble. (3) Apply RiteWeight until convergence to generated an estimated equilibrium distribution. Thus, the pipeline is applied independently to the output from each AI tool, although a common PC space is used to enable facile comparison of ensembles.

### System: Adenylate kinase

Human adenylate kinase (AK) exhibits substantial structural variability, most notably between open and closed conformations. Although both apo and liganded forms may populate each of these states to some extent, the relative populations under different conditions remain unknown. In this study, we focus on the wild-type protein. As the AI tools are trained on both apo and liganded structures without distinction, it is unclear whether the ensembles they generate represent the apo state, a liganded state, or some unphysical mixture

### AI ensemble generation

The AI ensembles were generated by two different AI tools: AFsample2 (v1.1) and ESMFlow (base MD simulations and PDB model 2024/02). The recommended settings of each AI tool were used to predict 10,000 structures of human AK based only on the residue sequence of human Adenylate kinase. By default, BioEmu filters unphysical structures (steric clashes or chain discontinuities), resulting in a slightly lower number of final structures.

For further analysis, all structures were aligned to a joint reference structure, and a principal component analysis was performed on the C*α* Cartesian coordinates of the aligned structures.

### Preparation of downsampled ensembles from full AI-generated ensembles

All AI tools were configured to generate 10,000 structural models. To identify an optimal structure for alignment, the resulting ensembles were first clustered using DBSCAN^50^ applied to pairwise distance matrices of backbone C*α* atoms, as implemented in the CPPTRAJ program of the Amber software package. Optimal DBSCAN parameters were determined using the k-distance (“elbow”) method, evaluated over k values ranging from 2 to 10. When multiple clusters were identified, the medoid of the largest cluster was selected as the representative structure for alignment; secondary clusters, when present, each comprised less than 1% of the dataset. Principal component (PC) analysis was then performed on the aligned C*α* Cartesian coordinates. The AFSample2 and ESMFlow ensembles were clustered jointly to generate a unified reference structure and PC space.

After projecting each AI-generated ensemble onto the first two PC’s, the space was partitioned into 10 equal-width bins along each component between the minimum and maximum values rounded to the nearest tenth. A downsampled ensemble was then generated by randomly selecting one structure from each occupied bin. Each selected structure was assigned a weight corresponding to the number of structures within its bin. Sidechains and any missing atoms were subsequently reconstructed for the downsampled ensemble, where necessary.

The resulting set of 20-80 structures was then minimized and equilibrated using the Amber ff14SBonlysc force field with the generalized Born implicit solvent model, GB-Neck2 (igb=8)^51,52^ at 298K and collision frequency (γ) of 80 ps^-1^. Each structure was subjected to 1000 steps of unrestrained energy minimization (500 steps of steepest descent followed by 500 steps of conjugate gradient), 50 ps of backbone-restrained NVT equilibration (1 kcal mol^-1^A^-2^), and 1 ps of unrestrained NVT equilibration.

### Weighted ensemble simulations

The weighted ensemble (WE) simulations were initiated from the minimized, equilibrated downsampled ensembles, with initial weights assigned according to the structure counts within each bin. The bin boundaries used in the WE simulations were identical to those used for downsampling the AI-generated ensembles, with the exception that 5 additional equally spaced bins were appended to each end of the two PC dimensions, resulting in a total of 400 WE bins. WE simulations were performed with a resampling time interval (τ) of 10 ps and a target of 8 trajectory segments per bin. Each simulation was run for one wall-clock day on 8 NVIDIA A100 GPUs and subsequently extended for an additional six days to improve sampling. Molecular dynamics (MD) simulation parameters were identical to those used at the final unrestrained equilibration step described above.

### RiteWeight

The RiteWeight algorithm^43^ was used to directly approximate equilibrium from the WE simulation data. RiteWeight employed a joint principal component space calculated during the initial analysis of the AI ensembles with a variance cut-off of 85%. Trajectory segments analyzed by the algorithm consist of the first and last structures of all WE iterations, corresponding to a time step τ = 10 ps. RiteWeight was applied using 2 clusters and 5000 iterations with an added smoothing of 1% to ensure convergence. In smoothed RiteWeight, the updated weight for a segment is given by the “raw” weight of the original algorithm^43^ averaged at the indicated percent with the cluster average weight. The initial weights were provided by the WE distribution.

## Results

The results for adenylate kinase (**Fig. 2**) suggest the pipeline is highly successful in this case. The initial AI ensembles, when projected onto PC1, are highly distinct, varying among mostly open, mostly closed, and bimodal distributions. WE simulations causes all the ensembles to relax toward more open conformations, and RiteWeight analysis yields final distributions that are similar to one another and significantly favor open structures. This finding is consistent with single-molecule FRET experiments.^53^

**Fig. 2.**
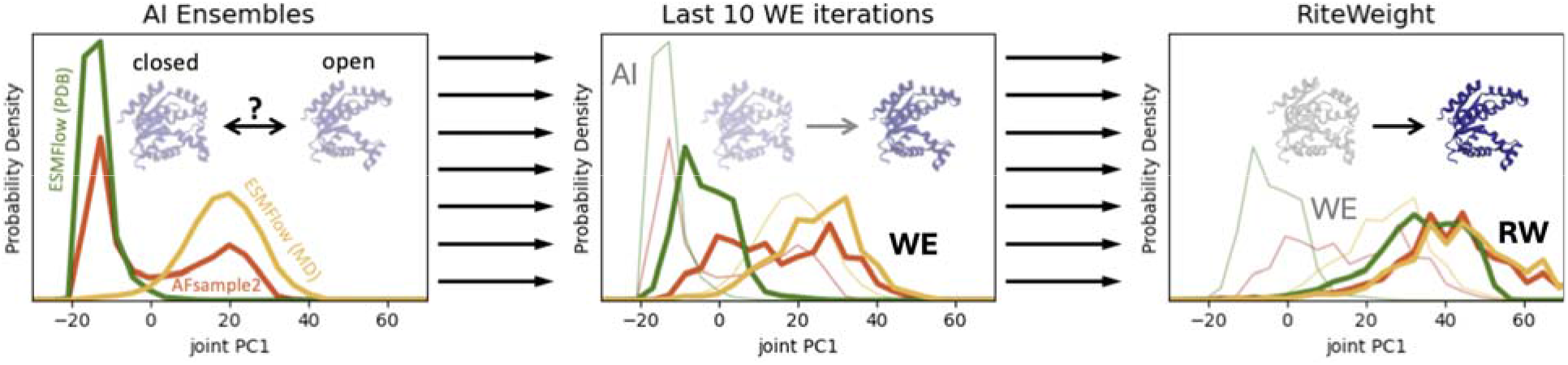
From highly distinct AI ensembles to approximated equilibrium distributions. Application of the AI-WE-RW pipeline to unliganded adenylate kinase. Left: Initial distributions from AI tools (thick lines) are highly dis-similar. Middle: WE simulation leads to noticeable relaxation toward larger values of PC1, corresponding to greater opening of the protein. WE distributions (thick lines) are averages over the final 10 iterations. Right: RiteWeight separately reweights each ensemble (thick lines) into a unimodal distribution favoring the open conformations.

## Concluding Discussion

We have described highly encouraging data from initial application of a set of two existing physics-based tools which markedly improve protein structural ensembles generated by AI tools. Weighted ensemble (WE) simulation, as implemented in the WESTPA package,^38,39^ simulates short-timescale relaxation from the initial AI-generated ensembles. In turn, WE data is used to directly approximate equilibrium via the recently introduced randomized iterative reweighting (RiteWeight) algorithm.^43^

The current data were obtained without significant optimization of the WE and RiteWeight protocols. WE simulation employed simple bins based on the first two principal components (PCs), and RiteWeight likewise was run in the space of PCs. In using RiteWeight, the smoothing protocol described above was vital to obtaining convergence without significant noise. Optimization of WE bins based on recent proposals,^54-56^ may improve WE data quality and thus RiteWeight convergence.

We recognize that the current simple protocol based on PCs might fall short for the most challenging systems. Because RiteWeight leverages local transition information in trajectories, it cannot succeed if WE trajectories fall into disjoint spaces without transitions between key states. For example, if AI ensembles exhibit well-separated conformational states in a complex system, it may be difficult to determine WE bins that enable bi-directional transitions between the states. However, a wide range of strategies have been proposed to improve WE convergence, progress coordinates, and bins.^54,56-64^

On the whole, we believe our initial findings hold promise for the near-term computational generation of equilibrium distributions consistent with a force field. As a consequence, with the apparently strong dependence of AI models on their training data, a pipeline like the one described here might provide valuable training data to the next iteration of AI models.

## Acknowledgements

LTC and DMZ gratefully acknowledge support from NIH Grant GM115805.

## Notes

### Competing Interest Statement

The authors have declared no competing interest.

## References

1 Lindorff-Larsen, K., Piana, S., Dror, R. O. & Shaw, D. E. How fast-folding proteins fold. Science 334, 517–520 (2011). 10.1126/science.1208351

2 Pande, V. S. Understanding protein folding using Markov state models. Adv Exp Med Biol 797, 101–106 (2014). 10.1007/978-94-007-7606-7_8

3 Marx, A., Hoenger, A. & Mandelkow, E. Structures of kinesin motor proteins. Cell Motil Cytoskeleton 66, 958–966 (2009). 10.1002/cm.20392

4 Yildiz, A. Mechanism and regulation of kinesin motors. Nat Rev Mol Cell Biol 26, 86–103 (2025). 10.1038/s41580-024-00780-6

5 Han, L. et al. Structure and mechanism of the SGLT family of glucose transporters. Nature 601, 274–279 (2022). 10.1038/s41586-021-04211-w

6 Davies, J. S., Currie, M. J., Dobson, R. C. J., Horne, C. R. & North, R. A. TRAPs: the ‘elevator-with-an-operator’ mechanism. Trends Biochem Sci 49, 134–144 (2024). 10.1016/j.tibs.2023.11.006

7 Song, H. D. & Zhu, F. Conformational dynamics of a ligand-free adenylate kinase. PLoS One 8, e68023 (2013). 10.1371/journal.pone.0068023

8 Arora, K. & Brooks, C. L., 3rd. Large-scale allosteric conformational transitions of adenylate kinase appear to involve a population-shift mechanism. Proc Natl Acad Sci U S A 104, 18496–18501 (2007). 10.1073/pnas.0706443104

9 Stiller, J. B. et al. Structure determination of high-energy states in a dynamic protein ensemble. Nature 603, 528–535 (2022). 10.1038/s41586-022-04468-9

10 Aranganathan, A., Gu, X., Wang, D., Vani, B. P. & Tiwary, P. Modeling Boltzmann-weighted structural ensembles of proteins using artificial intelligence–based methods. Current opinion in structural biology 91, 103000 (2025).

11 Amaro, R. E. et al. Ensemble Docking in Drug Discovery. Biophys J 114, 2271–2278 (2018). 10.1016/j.bpj.2018.02.038

12 Acharya, A. et al. Supercomputer-Based Ensemble Docking Drug Discovery Pipeline with Application to Covid-19. J Chem Inf Model 60, 5832–5852 (2020). 10.1021/acs.jcim.0c01010

13 Weikl, T. R. & Paul, F. Conformational selection in protein binding and function. Protein Sci 23, 1508–1518 (2014). 10.1002/pro.2539

14 Nussinov, R. Introduction to Protein Ensembles and Allostery. Chem Rev 116, 6263–6266 (2016). 10.1021/acs.chemrev.6b00283

15 Gianni, S., Dogan, J. & Jemth, P. Distinguishing induced fit from conformational selection. Biophys Chem 189, 33–39 (2014). 10.1016/j.bpc.2014.03.003

16 Broom, A. et al. Ensemble-based enzyme design can recapitulate the effects of laboratory directed evolution in silico. Nature Communications 11, 4808 (2020). 10.1038/s41467-020-18619-x

17 Crean, R. M., Gardner, J. M. & Kamerlin, S. C. L. Harnessing Conformational Plasticity to Generate Designer Enzymes. J Am Chem Soc 142, 11324–11342 (2020). 10.1021/jacs.0c04924

18 Hadzipasic, A. et al. Ancient origins of allosteric activation in a Ser-Thr kinase. Science 367, 912–917 (2020).

19 Oleinikovas, V., Saladino, G., Cossins, B. P. & Gervasio, F. L. Understanding cryptic pocket formation in protein targets by enhanced sampling simulations. Journal of the American Chemical Society 138, 14257–14263 (2016).

20 Nussinov, R., Liu, Y., Zhang, W. & Jang, H. Cell phenotypes can be predicted from propensities of protein conformations. Current opinion in structural biology 83, 102722 (2023).

21 Spinetti, E. et al. A Comparative Molecular Dynamics Study of Selected Point Mutations in the Shwachman-Bodian-Diamond Syndrome Protein SBDS. Int J Mol Sci 23 (2022). 10.3390/ijms23147938

22 el-Bastawissy, E., Knaggs, M. H. & Gilbert, I. H. Molecular dynamics simulations of wild-type and point mutation human prion protein at normal and elevated temperature. J Mol Graph Model 20, 145–154 (2001). 10.1016/s1093-3263(01)00113-9

23 Martinez-Zacarias, A. C., Lopez-Perez, E. & Alas-Guardado, S. J. Effect of the Lys62Ala Mutation on the Thermal Stability of BstHPr Protein by Molecular Dynamics. Int J Mol Sci 25 (2024). 10.3390/ijms25126316

24 Jumper, J. et al. Highly accurate protein structure prediction with AlphaFold. Nature 596, 583–589 (2021). 10.1038/s41586-021-03819-2

25 Lin, Z. et al. Language models of protein sequences at the scale of evolution enable accurate structure prediction. BioRxiv 2022, 500902 (2022).

26 Baek, M. et al. Accurate prediction of protein structures and interactions using a three-track neural network. Science 373, 871–876 (2021).

27 Kalakoti, Y. & Wallner, B. AFsample2 predicts multiple conformations and ensembles with AlphaFold2. Commun Biol 8, 373 (2025). 10.1038/s42003-025-07791-9

28 Jing, B., Berger, B. & Jaakkola, T. AlphaFold meets flow matching for generating protein ensembles. arXiv preprint arXiv:2402.04845 (2024).

29 Bhakat, S. & Strauch, E.-M. Accelerated sampling of protein dynamics using BioEmu augmented molecular simulation. bioRxiv, 2026.2001.2007.698041 (2026). 10.64898/2026.01.07.698041

30 Meller, A., Bhakat, S., Solieva, S. & Bowman, G. R. Accelerating Cryptic Pocket Discovery Using AlphaFold. Journal of Chemical Theory and Computation 19, 4355–4363 (2023). 10.1021/acs.jctc.2c01189

31 Vats, S., Bobrovs, R., Söderhjelm, P. & Bhakat, S. AlphaFold-SFA: Accelerated sampling of cryptic pocket opening, protein-ligand binding and allostery by AlphaFold, slow feature analysis and metadynamics. PLOS ONE 19, e0307226 (2024). 10.1371/journal.pone.0307226

32 Richman, D. D., Karaguesian, J.Suomivuori, C.-M. & Dror, R. O. Unlocking hidden biomolecular conformational landscapes in diffusion models at inference time. arXiv preprint arXiv:2512.03312 (2025).

33 Mehdi, S., Smith, Z., Herron, L., Zou, Z. & Tiwary, P. Enhanced sampling with machine learning. Annual Review of Physical Chemistry 75, 347–370 (2024).

34 Ribeiro, J. M. L., Bravo, P., Wang, Y. & Tiwary, P. Reweighted autoencoded variational Bayes for enhanced sampling (RAVE). J Chem Phys 149, 072301 (2018). 10.1063/1.5025487

35 Vani, B. P., Aranganathan, A., Wang, D. & Tiwary, P. AlphaFold2-RAVE: From Sequence to Boltzmann Ranking. J Chem Theory Comput 19, 4351–4354 (2023). 10.1021/acs.jctc.3c00290

36 Megías, A. et al. Iterative variational learning of committor-consistent transition pathways using artificial neural networks. Nature Computational Science, 1–11 (2025).

37 Tang, C. et al. Breaking the Timescale Barrier: Generative Discovery of Conformational Free-Energy Landscapes and Transition Pathways. arXiv preprint arXiv:2510.24979 (2025).

38 Zwier, M. C. et al. WESTPA: An interoperable, highly scalable software package for weighted ensemble simulation and analysis. Journal of chemical theory and computation 11, 800–809 (2015).

39 Russo, J. D. et al. WESTPA 2.0: High-performance upgrades for weighted ensemble simulations and analysis of longer-timescale applications. Journal of Chemical Theory and Computation 18, 638–649 (2022).

40 Zhang, B. W., Jasnow, D. & Zuckerman, D. M. The “weighted ensemble” path sampling method is statistically exact for a broad class of stochastic processes and binning procedures. The Journal of chemical physics 132 (2010).

41 Zuckerman, D. M. & Chong, L. T. Weighted ensemble simulation: review of methodology, applications, and software. Annual review of biophysics 46, 43–57 (2017).

42 Chong, L. T. & Zuckerman, D. M. Weighted Ensemble Simulation: Advances in methods, software, and applications. Wiley Interdisciplinary Reviews: Computational Molecular Science 15, e70055 (2025).

43 Kania, S., Webber, R. J., simpson, G., Aristoff, D. & Zuckerman, D. M. RiteWeight: Randomized Iterative Trajectory Reweighting for Steady-State Distributions Without Discretization Error. Proceedings of the National Academy of Sciences, Accepted (2026). 10.48550/arXiv.2401.05597

44 Bacci, M., Caflisch, A. & Vitalis, A. On the removal of initial state bias from simulation data. The Journal of chemical physics 150 (2019).

45 Bowman, G. R., Beauchamp, K. A., Boxer, G. & Pande, V. S. Progress and challenges in the automated construction of Markov state models for full protein systems. The Journal of chemical physics 131 (2009).

46 Eyal, E., Gerzon, S., Potapov, V., Edelman, M. & Sobolev, V. The limit of accuracy of protein modeling: influence of crystal packing on protein structure. Journal of molecular biology 351, 431–442 (2005).

47 Kleywegt, G. J. Experimental assessment of differences between related protein crystal structures. Biological Crystallography 55, 1878–1884 (1999).

48 Billeter, M. Comparison of protein structures determined by NMR in solution and by X-ray diffraction in single crystals. Quarterly reviews of biophysics 25, 325–377 (1992).

49 Spronk, C. A., Nabuurs, S. B., Krieger, E., Vriend, G. & Vuister, G. W. Validation of protein structures derived by NMR spectroscopy. Progress in Nuclear Magnetic Resonance Spectroscopy 45, 315–337 (2004).

50 Ester, M., Kriegel, H.-P., Sander, J. & Xu, X. in kdd. 226–231.

51 Nguyen, H., Roe, D. R. & Simmerling, C. Improved generalized born solvent model parameters for protein simulations. Journal of chemical theory and computation 9, 2020–2034 (2013).

52 Nguyen, H., Perez, A., Bermeo, S. & Simmerling, C. Refinement of generalized born implicit solvation parameters for nucleic acids and their complexes with proteins. Journal of chemical theory and computation 11, 3714–3728 (2015).

53 Aviram, H. Y. et al. Direct observation of ultrafast large-scale dynamics of an enzyme under turnover conditions. Proceedings of the National Academy of Sciences 115, 3243–3248 (2018).

54 Mitra, S., Biswas, R. & Chakrabarty, S. WeTICA: A directed search weighted ensemble based enhanced sampling method to estimate rare event kinetics in a reduced dimensional space. The Journal of Chemical Physics 162 (2025).

55 Ryu, W. H. et al. Reducing Weighted Ensemble Variance With Optimal Trajectory Management. Journal of Chemical Physics, Accepted (2025).

56 Torrillo, P. A., Bogetti, A. T. & Chong, L. T. A minimal, adaptive binning scheme for weighted ensemble simulations. The Journal of Physical Chemistry A 125, 1642–1649 (2021).

57 Prabhakar, P. R., Ray, D. & Andricioaei, I. Discriminant analysis optimizes progress coordinate in weighted ensemble simulations of rare event kinetics. The Journal of Chemical Physics 163 (2025).

58 Adelman, J. L. & Grabe, M. Simulating rare events using a weighted ensemble-based string method. The Journal of chemical physics 138 (2013).

59 Ahn, S.-H., Ojha, A. A., Amaro, R. E. & McCammon, J. A. Gaussian-accelerated molecular dynamics with the weighted ensemble method: A hybrid method improves thermodynamic and kinetic sampling. Journal of chemical theory and computation 17, 7938–7951 (2021).

60 Dickson, A. & Brooks III, C. L. WExplore: hierarchical exploration of high-dimensional spaces using the weighted ensemble algorithm. The Journal of Physical Chemistry B 118, 3532–3542 (2014).

61 Leung, J. M. et al. Unsupervised Learning of Progress Coordinates during Weighted Ensemble Simulations: Application to NTL9 Protein Folding. Journal of Chemical Theory and Computation (2025).

62 Ojha, A. A., Thakur, S.Ahn, S.-H. & Amaro, R. E. DeepWEST: Deep learning of kinetic models with the Weighted Ensemble Simulation Toolkit for enhanced sampling. Journal of chemical theory and computation 19, 1342–1359 (2023).

63 Wang, D. & Tiwary, P. Augmenting Human Expertise in Weighted Ensemble Simulations through Deep Learning-Based Information Bottleneck. Journal of chemical theory and computation 20, 10371–10383 (2024).

64 Donyapour, N., Roussey, N. M. & Dickson, A. REVO: Resampling of ensembles by variation optimization. The Journal of chemical physics 150 (2019).

